# The far extracellular CUB domain of the adhesion GPCR ADGRG6/GPR126 is a key regulator of receptor signaling

**DOI:** 10.1101/2024.02.16.580607

**Authors:** Sumit J. Bandekar, Ethan E. Dintzner, Katherine Leon, Szymon P. Kordon, Tomasz Slezak, Kristina Cechova, Reza Vafabakhsh, Demet Araç

## Abstract

Adhesion G protein-coupled receptors (aGPCRs) transduce extracellular adhesion events into cytoplasmic signaling pathways. ADGRG6/GPR126 is an aGPCR critical for axon myelination, heart development and ear development; ADGRG6 is also associated with developmental diseases and cancers. ADGRG6 has a large, alternatively spliced, five-domain extracellular region (ECR) that samples different conformations and is essential for receptor function in vivo. However, the mechanistic details of how the ECR regulates signaling are unclear. Herein, we studied the conformational dynamics of the conserved CUB domain which is located at the distal N-terminus of the ADGRG6 ECR and is deleted in an alternatively spliced isoform (ΔCUB). We show that the ΔCUB isoform has decreased signaling and is insensitive to inclusion of an activating splice insertion (+ss). Molecular dynamics simulations suggest that the CUB domain is involved in interdomain contacts to maintain a compact ECR conformation. A cancer-associated CUB domain mutant, C94Y, drastically perturbs the ECR conformation and results in elevated signaling, whereas another CUB mutant located near a conserved Ca^2+^-binding site, Y96A, decreases signaling. Our results suggest an ECR-mediated mechanism for ADGRG6 regulation in which the CUB domain instructs conformational changes within the ECR to regulate receptor signaling.

## Introduction

Multicellular organisms employ cell-surface receptors to coordinate the intercellular events that are necessary for developmental and regulatory processes. Adhesion G protein-coupled receptors (aGPCRs) represent a diverse and understudied family of cell-surface receptors that couple cellular adhesion to signaling (1). aGPCRs have critical roles in synapse formation, myelination of the nervous system, angiogenesis, neutrophil activation, embryogenesis, and other biological functions (1–4). Additionally, aGPCRs are strongly implicated in developmental diseases, neurological diseases, and cancer (1, 5, 6). Despite their myriad of functions and roles in disease, the molecular mechanisms underlying the function of many aGPCRs remain poorly understood.

aGPCRs contain characteristically large, multidomain extracellular regions (ECRs) with adhesion domains that bind to ligands (7–15). Generally, the ECR contains a conserved GPCR autoproteolysis inducing (GAIN) domain proximal to the seven transmembrane (7TM) region; this GAIN domain is autoproteolyzed in many aGPCRs. According to the tethered agonist (TA) model of aGPCR activation, removal of the ECR from the 7TM reveals a TA peptide which then activates the receptor (1, 16). A growing pool of evidence shows that the ECR can also directly modulate 7TM signaling in a TA-independent manner (1, 11, 12, 14, 17–19). In this second model, direct allosteric communication between ECR and the 7TM is important for the regulation of receptor activity (1, 19). ECR conformation can be affected through the binding of a ligand or by alternative splicing (14, 20) and these conformational changes can be communicated to the 7TM to alter signaling (19). In further support of this idea, roughly half of aGPCRs lack TA-dependent signaling in vitro (21). For many aGPCRs, the mechanistic basis by which ECR conformation regulates receptor activity is incompletely understood.

ADGRG6/GPR126 is an evolutionarily conserved aGPCR critical for peripheral nervous system (PNS) myelination, skeletal development, angiogenesis, and heart development (22–26). The defining characteristic of ADGRG6 is a large 95 kilodalton (kDa) ECR consisting of the Complement C1r/C1s, Uegf, and Bmp1 (CUB), Pentraxin (PTX), sperm protein, enterokinase and agrin (SEA), Hormone Receptor (HormR), and GAIN domains (Fig. 1A) (14). During PNS myelination, the ECR of ADGRG6 and downstream ADGRG6 signaling is necessary for axon sorting (27–29). Furthermore, the ECR of ADGRG6 is required for its function in heart development (26). Mutations in the ADGRG6 ECR result in loss of PNS myelination and are causative in arthrogryposis multiplex congenita (AMC) (30). Mutations in the ECR of ADGRG6 could contribute to adolescent idiopathic scoliosis (AIS) disease progression (14, 31). Downstream of the receptor, the roles of ADGRG6 in development have been tied to its ability to initiate signaling through the second messenger cyclic adenosine monophosphate (cAMP) (27–29).

**Figure 1.**
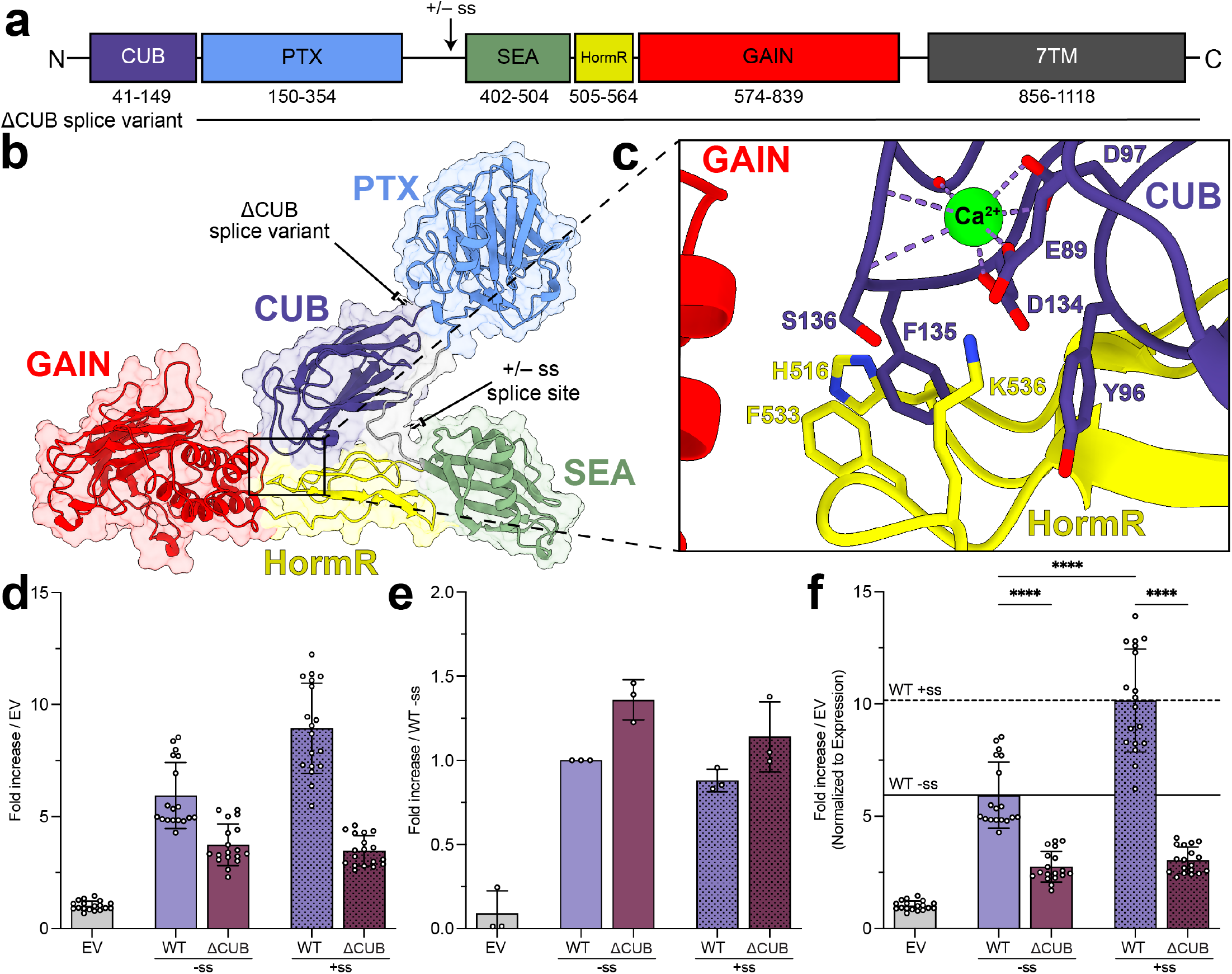
Deletion of the CUB domain decreases ADGRG6-dependent cAMP signaling. **(a):** Linear domain diagram for ADGRG6. The ΔCUB splice variant is depicted as a horizontal black line to show its domain boundaries. The position of the splice site between PTX and SEA is designated by an arrow. **(b):** ADGRG6 –ss ECR crystal structure (PDB 6V55) depicted using cartoon along with surface representation. **(c):** Close-up of the CUB/HormR interface from the crystal structure with key residues involved in the interaction shown as sticks. The bound Ca^2+^ ion is shown as a green sphere with coordination shown as dotted lines. **(d):** Basal cAMP signaling of ADGRG6 constructs shown as fold change over empty vector (EV). Mutants are grouped by absence (–ss) or presence (+ss) of the ECR splice insert. N=3 independent experiments were performed with six technical replicates each. **(e):** Relative cell surface expression levels for ADGRG6 constructs shown as fold increase over WT –ss. See Fig. S1 for raw flow cytometry data. N=3 independent experiments were performed. **(f):** Basal cAMP signaling shown as fold change over empty vector (EV) normalized to cell-surface expression. Mutants are grouped by absence (–ss) or presence (+ss) of the ECR splice insert. Horizontal lines show the mean normalized signaling levels for WT constructs. **** – p<0.0001; one–way ANOVA test with Tukey’s correction for multiple comparisons. Each condition was compared to every other condition, but not all comparisons are shown for clarity. Bar graphs represent mean values with error bars showing standard deviation.

The conformation of the ADGRG6 ECR is associated with its downstream signaling level (14). Previous work suggested that a series of open-like conformations where the N-terminal CUB domain extends away from the other ECR domains drives a higher level of signaling activity. In contrast, a closed and compact ECR conformation formed by the CUB domain engaging the HormR domain, which was observed in a high-resolution crystal structure, corresponds to a lower signaling level (14). These conformations are regulated by alternative splicing, where the inclusion (+ss) of a 23-residue splice insert between the PTX and SEA domains (Fig. 1A) shifts the ECR towards the open-like conformation and higher signaling activity (14). The –ss splice variant is predominantly found in the closed conformation and has lower signaling activity. The key difference between these conformations is the location of the CUB domain relative to other ECR domains (14), suggesting its importance in receptor signaling.

The ADGRG6 CUB domain is the most conserved domain in the ECR and occupies the far N-terminal position, enabling it to reach farthest into the extracellular space and interact with ligands. During zebrafish development, transcripts are detected for alternatively spliced variants encoding ADGRG6 without its CUB domain (termed ΔCUB herein) (Fig. 1A) (26), suggesting that the CUB domain plays an important regulatory role during development. The most conserved portion of the ADGRG6 CUB domain is a Ca^2+^ binding site coordinating a single Ca^2+^ ion. The Ca^2+^ binding site on ADGRG6 constitutes a putative ligand binding surface as this region is used by other CUB domains to bind extracellular protein ligands (32–34). In the closed ECR conformation of ADGRG6 observed in its crystal structure (14), the CUB domain uses this Ca^2+^ binding surface to engage the HormR domain (Fig. 1B, C). Point mutations on and around the Ca^2+^ binding site lead to PNS myelination and ear development defects in zebrafish (14). Additionally, the binding of one putative ADGRG6 ligand, Collagen IV (CIV), has been confined to the CUB-PTX tandem (35). Taken together, ADGRG6 ECR conformation regulates signaling activity through a mechanism involving residues on the Ca^2+^ binding site within the CUB domain. However, the molecular details of this process remain unclear, precluding a comprehensive understanding of ADGRG6 regulation.

In this work, we employ a combination of signaling assays and microsecond-length molecular dynamics (MD) simulations to understand the mechanism by which CUB regulates the function of ADGRG6. We find that the ΔCUB splice variant results in lower cAMP signaling and this variant loses sensitivity to +ss. Using MD simulations, we show that the –ss ADGRG6 ECR adopts multiple closed conformations where the CUB Ca^2+^ binding site remains bound to HormR. In our MD simulations, we identified an interface formed between the PTX and SEA domains that, when mutated, lowers ADGRG6 signaling and abrogates sensitivity to +ss. The cancer associated mutant C94Y, which is on the CUB domain but distal from the Ca^2+^ binding site, displaces the CUB domain from HormR in MD, and has higher signaling activity than the WT receptor. Another cancer-associated mutant, Y96A, located on the Ca^2+^ binding surface, fails to access the higher signaling state of the +ss isoform. These results suggest that exposure of the CUB domain to solvent stimulates ADGRG6 signaling as long as the Ca^2+^ binding surface is not affected.

## Results

### The ΔCUB splice variant decreases ADGRG6 cAMP signaling

A developmental splice variant of ADGRG6 has been detected (26) which lacks the N-terminal CUB domain (ΔCUB, Fig. 1A). As several functions of ADGRG6 depend on its capacity to signal through the cAMP second messenger (27–29), we first assessed the ability of ADGRG6 ΔCUB to elevate intracellular cAMP levels. We employed a cAMP assay using HEK293 cells transiently transfected with zebrafish ADGRG6 (Fig. 1D) (14, 28). We tested all constructs for expression using flow cytometry against an N-terminal FLAG tag (Fig. 1E, Fig. S1A-D) and found that ΔCUB constructs expressed similarly to WT ADGRG6 constructs. Measured cAMP levels were normalized to the expression of each construct (Fig. 1F). As previously shown (14), WT +ss ADGRG6 elevates cAMP to a significantly higher level than the –ss receptor (Fig. 1F). However, we observed both –ss and +ss isoforms of ΔCUB exhibited significantly less cAMP elevation than their WT counterparts (Fig. 1F). Furthermore, the +ss splice site-dependent elevation in signaling observed with WT ADGRG6 was not observed in the ΔCUB variants. This loss of cAMP elevation was insensitive to pertussis toxin (PTx) addition (Fig. S1 E-H), ruling out the possibility that the ΔCUB splice variants could exhibit lower cAMP signaling due to enhanced Gα_i_ coupling. Additionally, WT and ΔCUB ADGRG6 were both insensitive to addition of reported ligands in our system (Fig. S1I, J), including the cellular prion protein (PrP^c^) (36) and Collagen IV (CIV) (35). ΔCUB ADGRG6 variants were less activated by the TA peptide added in trans vs. WT (Fig. S1K, L). These results suggest that the CUB domain is required for the signaling of ADGRG6 and that the ΔCUB splice variant likely plays a distinct developmental role from the WT receptor as its ability to elevate cAMP is diminished.

### Microsecond–timescale MD simulations of the ADGRG6 –ss ECR show sustained interdomain contacts involving the CUB, PTX, and SEA domains

As the conformational state of the ADGRG6 ECR is associated with signaling (14), we sought to understand the conformational dynamics of the ECR. While the structure of the closed/compact ECR conformation (Fig. 2A) has already been determined (14), the structure of the open-like and active conformation is not known. We crystallized the ADGRG6 +ss ECR at 3.7 Å resolution (Fig. S2, Supporting information, Table S1). The crystals contain density for the SEA/HormR/GAIN domains and several other indications that the solution is correct (Fig. S2, Supporting information). However, we do not observe any density for the CUB/PTX domains, suggesting that they may be flexible relative to SEA/HormR/GAIN (Fig. 2B). Since we could not observe the CUB domain in this structure, we utilized MD simulations to understand the conformational space accessed by to the ADGRG6 ECR.

**Figure 2.**
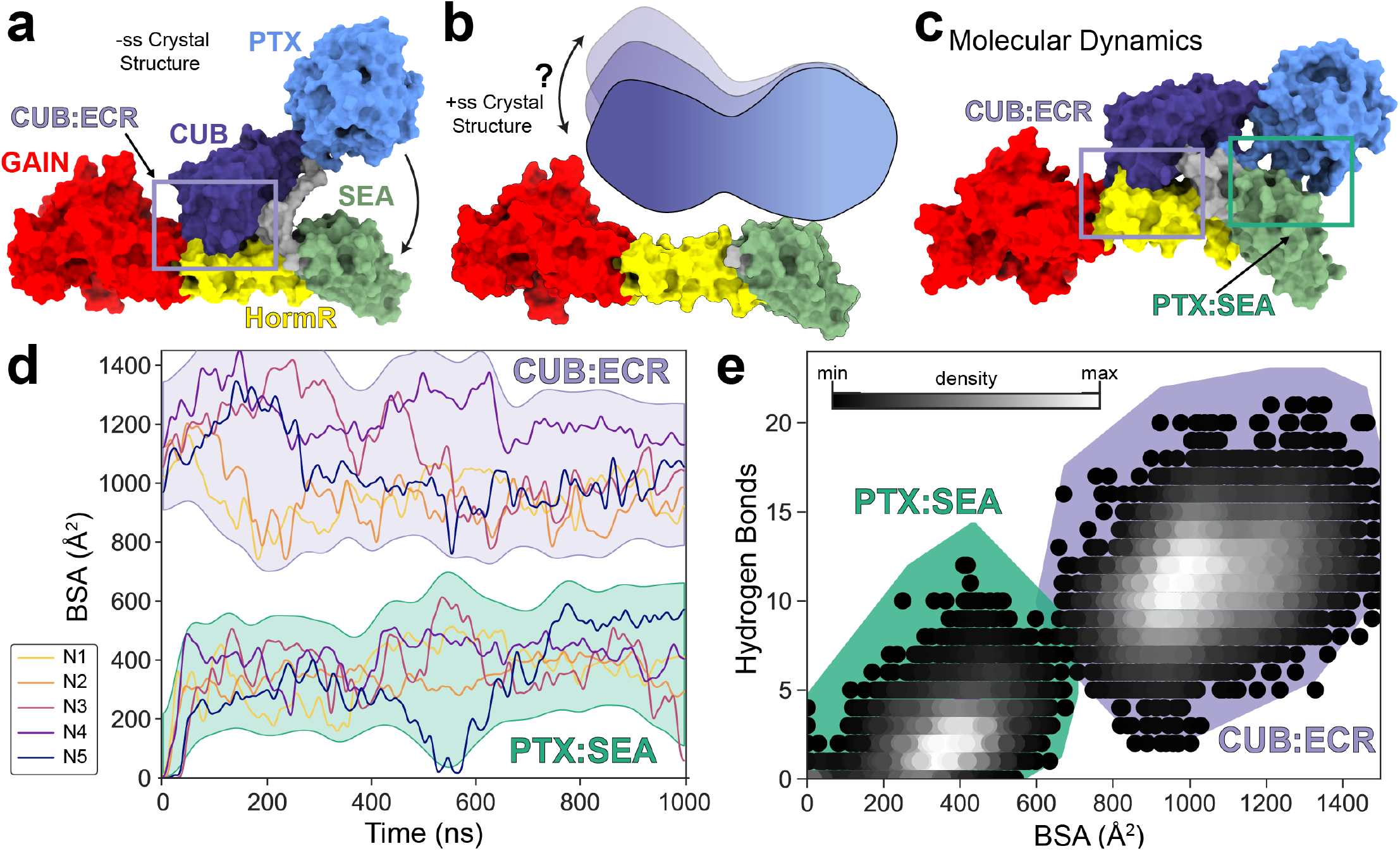
Molecular dynamics simulations of the ADGRG6 ECR show sustained interdomain contacts involving the CUB, PTX, and SEA domains. **(a):** Surface representation of -ss ADGRG6 from PDB: 6V55 with the CUB:ECR interface shown as a light purple rectangle. **(b):** The crystal structure of the +ss ADGRG6 ECR does not have the CUB and PTX domains resolved, suggesting they are flexible relative to the SEA/HormR/GAIN domains. See Fig. S2 for more information about the +ss ADGRG6 ECR crystal structure. **(c):** A representative frame from the MD simulations showing the conformational differences from the –ss crystal structure. The CUB:ECR and PTX:SEA interfaces are shown as light purple and light green rectangles, respectively. **(d):** Buried surface area (BSA) of CUB:ECR and PTX:SEA interfaces plotted over simulation time. Colored lines show each individual simulation, and the shaded areas represent the upper and lower bounds of all simulations. **(e):** The number of hydrogen bonds plotted versus the BSA of the described interfaces. The density (% of frames) of each position is plotted as a series of circles, with the most occupied regions of the plot in white, and the least in black.

We performed five one–microsecond all–atom MD simulations initialized from the crystal structure of the –ss ADGRG6 ECR (Fig. S3A–D, and Movie S1). Throughout these simulations, the ADGRG6 CUB domain remained associated with HormR. However, the observed ECR conformations were markedly different from the crystal structure (Fig. 2A vs. Fig. 2C). The CUB domain interacted with the ECR through a robust interface (CUB:ECR) involving HormR and the linker between PTX and SEA domains through several interactions present in the crystal structure, as well as several additional contacts (Fig. 2C, Fig. S3E). The CUB:ECR interface included both the Ca^2+^ binding site as well as other CUB domain surfaces. We also observed the PTX domain docking onto the SEA domain to form an interface (PTX:SEA), which remained intact for the remainder of the simulations. To quantify the extent of the CUB:ECR and PTX:SEA interactions, we performed buried surface area (BSA) and hydrogen bond calculations (Fig. 2D, E). Throughout all simulations, the CUB domain buried approximately 1000 Å^2^ from solvent and formed on average 10 hydrogen bonds with the rest of the ECR. The PTX:SEA interface buries approximately 350 Å^2^ from solvent and forms on average 3 interdomain hydrogen bonds. These results suggest that the CUB interaction with the ECR is mediated by a robust network of hydrophobic and hydrophilic contacts involving both the Ca^2+^ binding site and other CUB surfaces, and that the PTX domain could associate with the SEA domain in solution.

### ADGRG6 –ss adopts multiple conformations which are distinct from the crystal structure and involve large conformational changes of the CUB–PTX tandem

To quantify ADGRG6 ECR conformational dynamics, we performed principal component analysis (PCA) on our simulations. PCA simplifies the complex conformational landscape observed in our simulations into concerted motions called principal components (PCs). PCA identified two PCs which capture 76% of the conformational variance of our system (Fig. S3F–I). The first of these two PCs describes a 47º bend of the CUB, PTX, SEA, and HormR domains about the GAIN domain with the CUB–PTX tandem contributing the most inertia (Fig. 3A and Movie S2). The second PC is represented by an orthogonal 36º bending motion of CUB, PTX, SEA, and HormR domains about the GAIN domain with similar deviation of all domains from the average structure (Fig. 3B and Movie S3). During the simulations, a variety of regions in PC1-2 space are sampled, highlighting that ADGRG6 can adopt several metastable conformations (Fig. 3C). Hierarchical clusters were generated in PC1-2 latent space to classify the simulations, revealing clusters whose centroids are distinct conformations from the crystal structure (Fig. 3D) with Cα RMSDs of 18.6 Å, 32.8 Å, 21.4 Å, and 28.4 Å for centroids 1, 2, 3, and 4 respectively. These data show that the ADGRG6 –ss ECR adopts several conformational states largely through motions of the CUB-PTX tandem, but throughout all these states the CUB domain including its Ca^2+^ binding site remains docked to its binding site on HormR.

**Figure 3.**
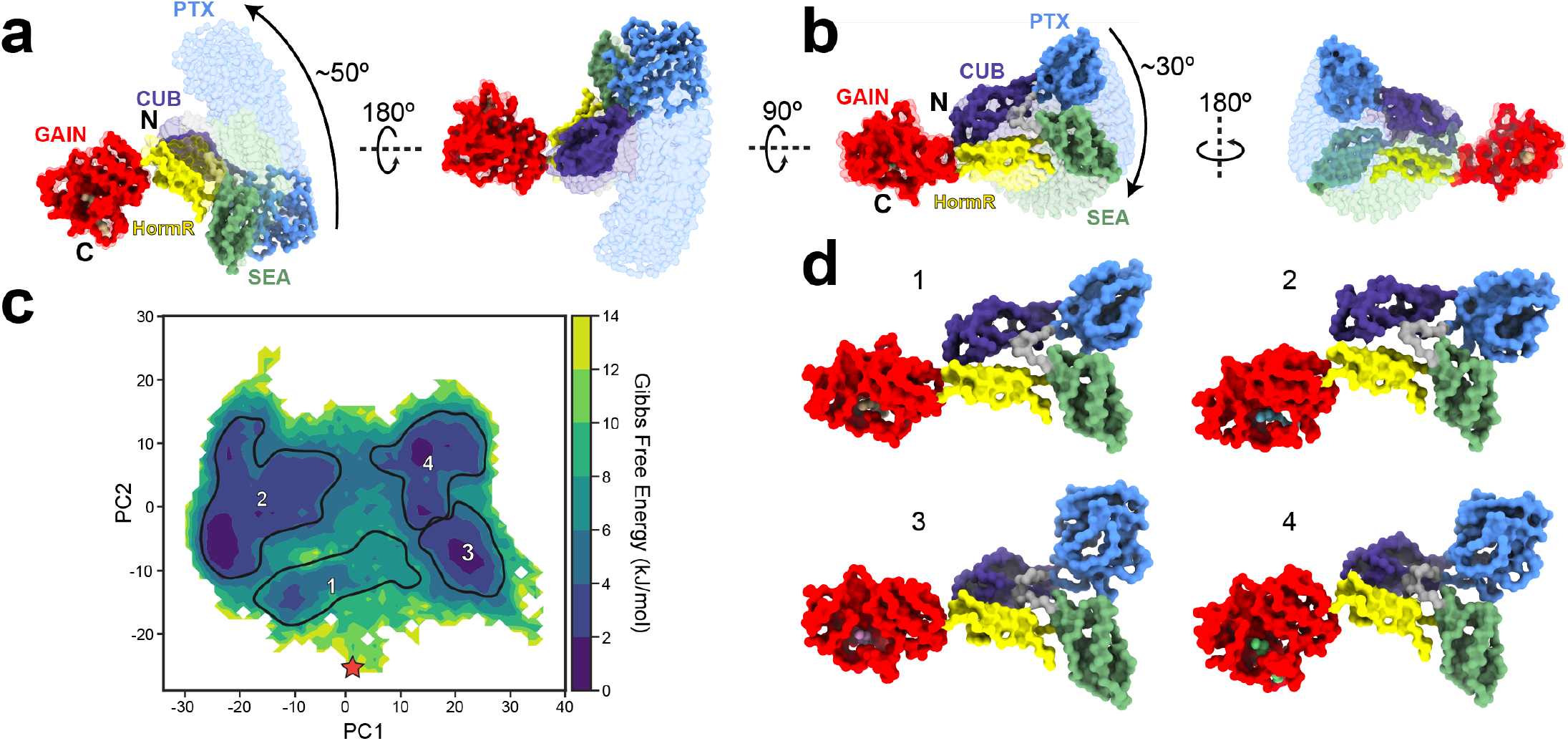
Principal component analysis reveals multiple conformations of ADGRG6 –ss ECR with the CUB domain putative ligand binding site docked onto HormR. Principal component analysis was performed on the α-carbon coordinates of N=5 concatenated WT ADGRG6 –ss ECR simulations. **(a):** Projection of the α-carbons along PC1 shows a 47º bend of the ECR about the GAIN domain. The initial (left) and final (right) frames of the projection are shown as opaque structures, and intermediates are displayed transparently. **(b):** Projection of the α-carbons along PC2 shows a 34º bend of the ECR about the GAIN domain orthogonal to PC1. Shown as in (a). **(c):** Free energy surface calculated in PC1-2 space. The red star represents the starting frame of the simulation. Black borders labeled 1-4 denote 75% boundaries of the probability mass function for hierarchical clusters on this free energy surface. The positions of these labels represent the coordinates of the centroids for the corresponding cluster. **(d):** The four centroids from panel (c) are shown as α-carbon surface representations to represent favored conformations across ADGRG6 –ss ECR simulation space.

### Residues in the PTX:SEA interface mediate ADGRG6 signaling levels

We analyzed the PTX:SEA interaction interface formed in our MD simulations and found that it was largely composed of residues K162-165 on the PTX domain, and a single face of the α2 helix on the SEA domain (Fig. 4 A, B). Some of these residues are mutated in human cancer patients, including S163 (T163A in Uterine Endometrioid |Carcinoma) and E482 (Q498E in Head and Neck Squamous Cell Carcinoma) (37). The residues in the PTX:SEA interface are variably conserved across animal species (Fig. 4C). We employed the ADGRG6 cAMP assay (Fig. 4D) and tested all constructs for expression using flow cytometry against an N-terminal FLAG tag (Fig. 4E, Fig. S4A, B). Measured cAMP levels were normalized to the expression of each construct (Fig. 4F). While –ss ADGRG6 PTX:SEA mutants displayed no statistical difference from WT, +ss ADGRG6 PTX:SEA mutants displayed lower cAMP signaling than WT, suggesting that they cannot access the open-like ECR conformations that correlate with higher signaling.

**Figure 4.**
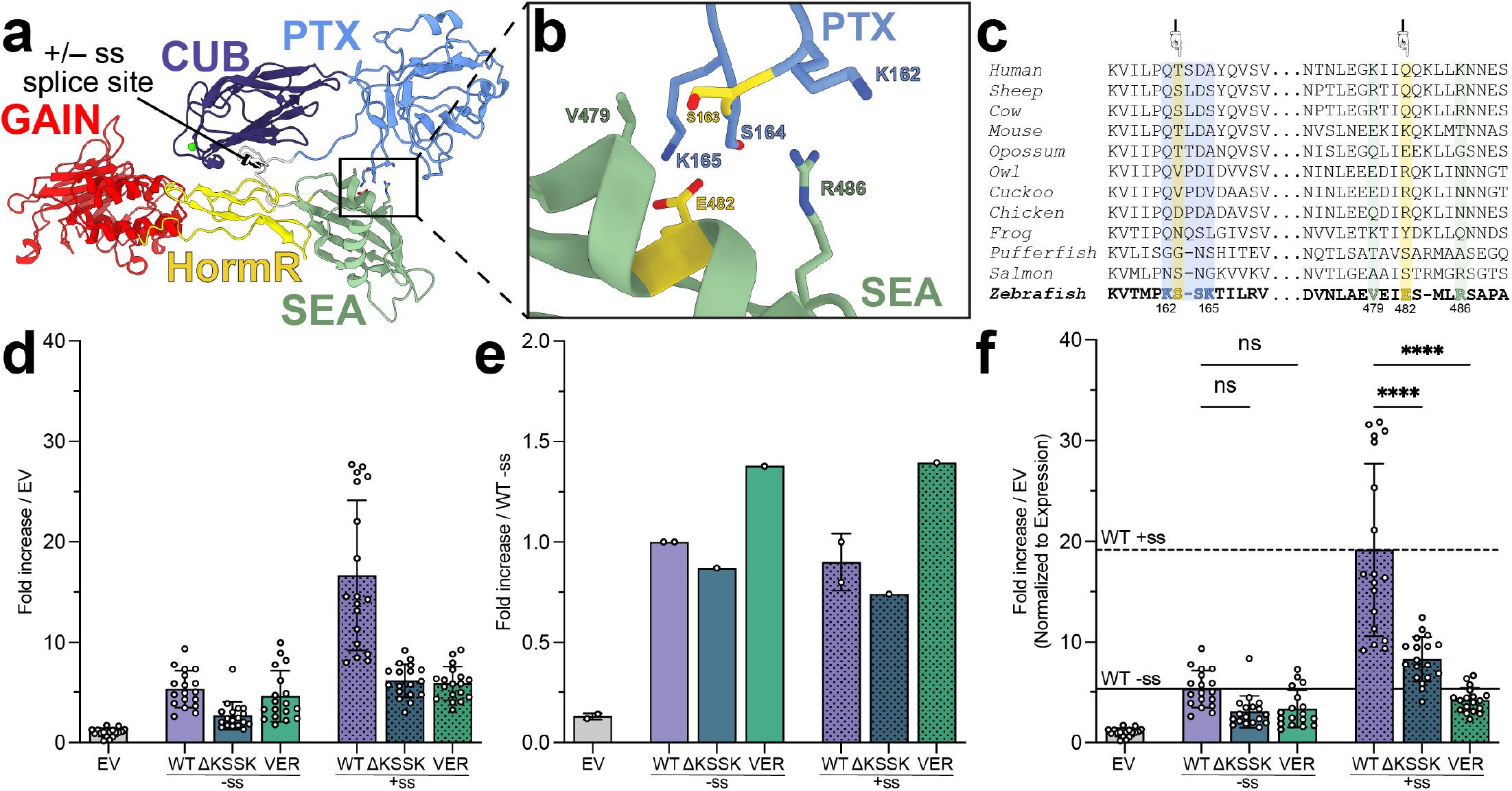
Mutation of the PTX:SEA interface decreases ADGRG6 signaling. **(a):** A representative frame from the MD simulations highlighting the interface between PTX and SEA domains. **(b):** Close-up of the PTX:SEA interface with key residues involved in the interaction shown as sticks. Residues which vary in human cancer (cBioPortal) are highlighted in yellow. **(c):** Sequence conservation of residues in the PTX:SEA interface is shown using ADGRG6 orthologs. Residues on the PTX domain are highlighted in silk blue, residues on the SEA domain are highlighted in light green, and residues which vary in cancer are highlighted in yellow and pointed to with arrows. **(d):** Basal cAMP signaling of ADGRG6 constructs shown as fold change over empty vector (EV). ΔKSSK refers to the ΔK162-K165 construct, and VER refers to the V479A/E482A/R486A construct. Mutants are grouped by absence (–ss) or presence (+ss) of the ECR splice insert. N=3 independent experiments were performed with six technical replicates each. **(e):** Relative cell surface expression levels for ADGRG6 constructs shown as fold increase over WT –ss. See Fig. S4 for raw flow cytometry data. N=1 independent experiments were performed. **(f):** Basal cAMP signaling shown as fold change over empty vector (EV) normalized to cell-surface expression. Mutants are grouped by absence (–ss) or presence (+ss) of the ECR splice insert. Horizontal lines show the mean normalized signaling levels for WT constructs. **** – p<0.0001; ns – not significant; one–way ANOVA test with Tukey’s correction for multiple comparisons. Each condition was compared to every other condition, but not all comparisons are shown for clarity. Bar graphs represent mean values with the standard deviation shown for error.

### A cancer-associated CUB mutation, C94Y, can occupy conformations which break the CUB:ECR interface and expose the Ca^2+^ binding site

The ADGRG6 CUB domain mediates the closed ECR conformation through an extensive contact interface involving not only the Ca^2+^ binding site, but also several distal residues. To investigate the effect of point mutations on the closed conformation, we simulated the C94Y mutant which is present in cancer patients (37). C94Y breaks a disulfide bond between C94 and C111 between CUB and HormR in a CUB domain loop distal to the Ca^2+^ binding site (Fig. 1C). We ran three one-microsecond simulations (Fig. 5, Fig. S5, and Movie S4) and observed the ECR settle into various energy minima (Fig. 5A). We also quantified the BSA of the CUB:ECR and PTX:SEA interfaces throughout the simulations (Fig. 5B). Strikingly, during the C94Y N=1 simulation, the CUB:ECR BSA dropped sharply and remained substantially lower than any other run performed (Fig. 5B). We performed PCA on a merged dataset including the C94Y N=1 and the five WT runs and found that considerable portions of the C94Y N=1 simulation occupy a unique region of the resulting PC1-2 latent space (Fig. 5C, red points Fig. S5E, and Movie S5, S6). Over the course of the simulation (Fig. 5D), the C94Y N=1 CUB domain shifted away from the HormR domain and ultimately dissociated from the ECR. The CUB domain then bound to a lateral face of HormR, leaving the Ca^2+^ binding site exposed to the solvent. This new, open-like, state is favored over the conformation where CUB is docked to HormR (ΔG = –2 kJ/mol) despite involving a loss of ∼500 Å^2^ in BSA between CUB and the ECR (Fig. 5A, B). These results show that the C94Y mutant can adopt a favorable open-like conformation unique from the WT ECR where the Ca^2+^ binding site is exposed to solvent.

**Figure 5.**
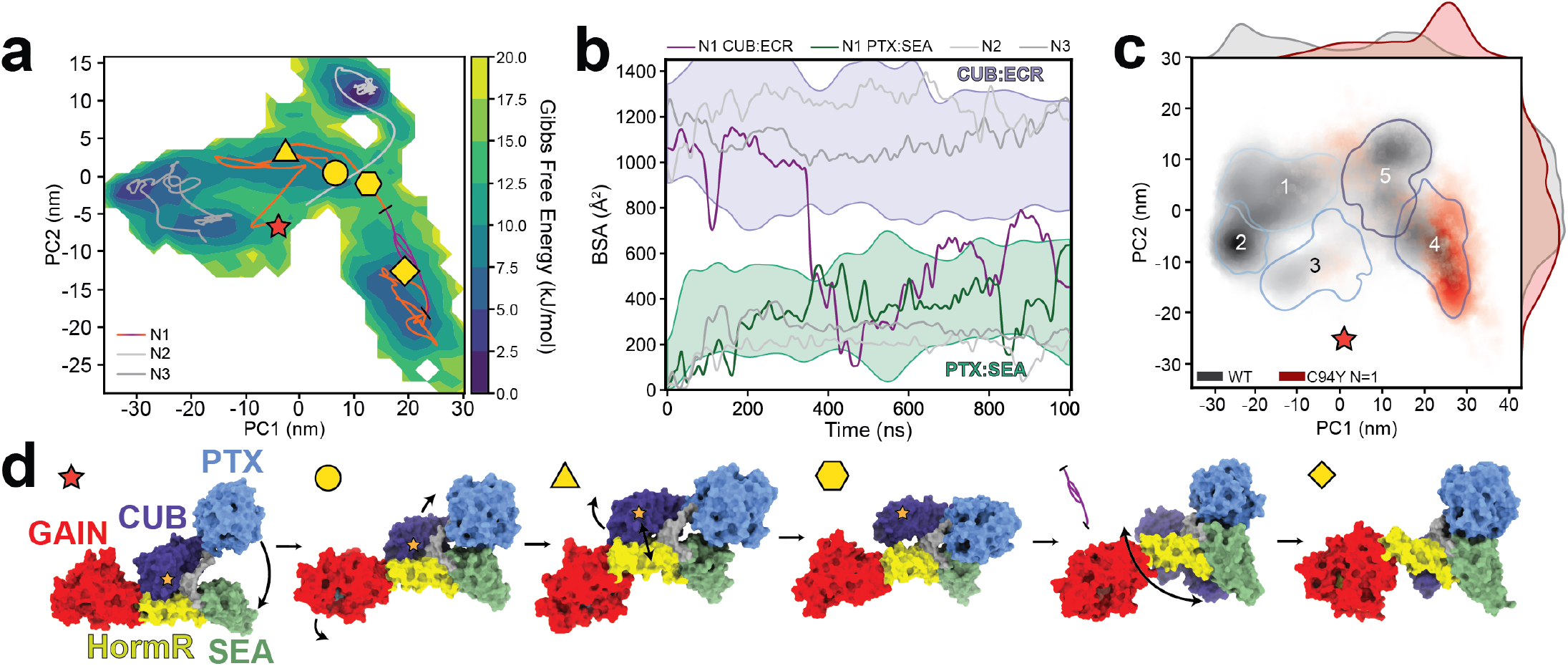
The ADGRG6 –ss ECR mutant C94Y can sample conformations which break the CUB:ECR interface and expose the CUB ligand binding site. **(a):** Free energy surface calculated in PC1-2 space. An orange line indicates the C94Y N=1 simulation trajectory, with different shapes corresponding to the structures shown in panel d. A purple line segment is used to denote a portion of the simulation where the CUB domain dissociates from HormR. The N=2 and N=3 trajectories are shown in shades of grey. **(b):** Buried surface area (BSA) of CUB:ECR and PTX:SEA interfaces over time. Dark purple and green lines represent the C94Y N=1 simulation CUB:ECR and PTX:SEA interfaces respectively, and the shaded intervals represent the bounds of the WT simulations for the corresponding interfaces. N=2 and N=3 are shown in shades of grey. **(c):** Comparison of the WT (greyscale) and C94Y N=1 simulations (red) in a combined essential space. Blue borders show 85% boundaries of the probability mass function for each hierarchical cluster. 1D histograms on the top and right of the square represent density of projections of each simulation along each PC. The red star is a projection of the starting point of the simulation onto this PC space. **(d):** Representative MD frames of C94Y –ss N=1 show conformational transitions throughout the simulation. The starting point of the simulation is represented as a red star and the location of C94Y is marked by a gold star. PTX docks onto SEA (yellow circle), and then CUB dissociates from HormR (yellow triangle, then yellow hexagon). After CUB samples several conformations (purple bounded line, represented as a series of CUB positions), it docks against the side of HormR with its ligand binding surface exposed (yellow diamond).

### Cancer-associated mutations in the Ca^2+^ binding surface affect receptor activity

We located several cancer-associated mutations in the CUB:ECR interface (Fig. 6A, B) (37). To understand how our cancer-associated mutants may affect ADGRG6 function, we employed the cAMP assay to test ADGRG6 signaling levels (Fig. 6C). We tested C94Y, which can induce an open-like ECR conformation despite its distal position from the Ca^2+^ binding surface. We also tested Y96A, a cancer-associated mutation on the Ca^2+^ binding surface which changes the highly conserved tyrosine that participates in ligand binding in several CUB domains (14, 32–34). We selected two further cancer-associated mutations in the interface between CUB and HormR, G372W and P520Q. We assessed the expression levels of ADGRG6 mutants with flow cytometry using an antibody directed against the N–terminal FLAG tag (Fig. 6D, Fig. S6A, B) and found that all mutants express to a similar level as ADGRG6 WT.

**Figure 6.**
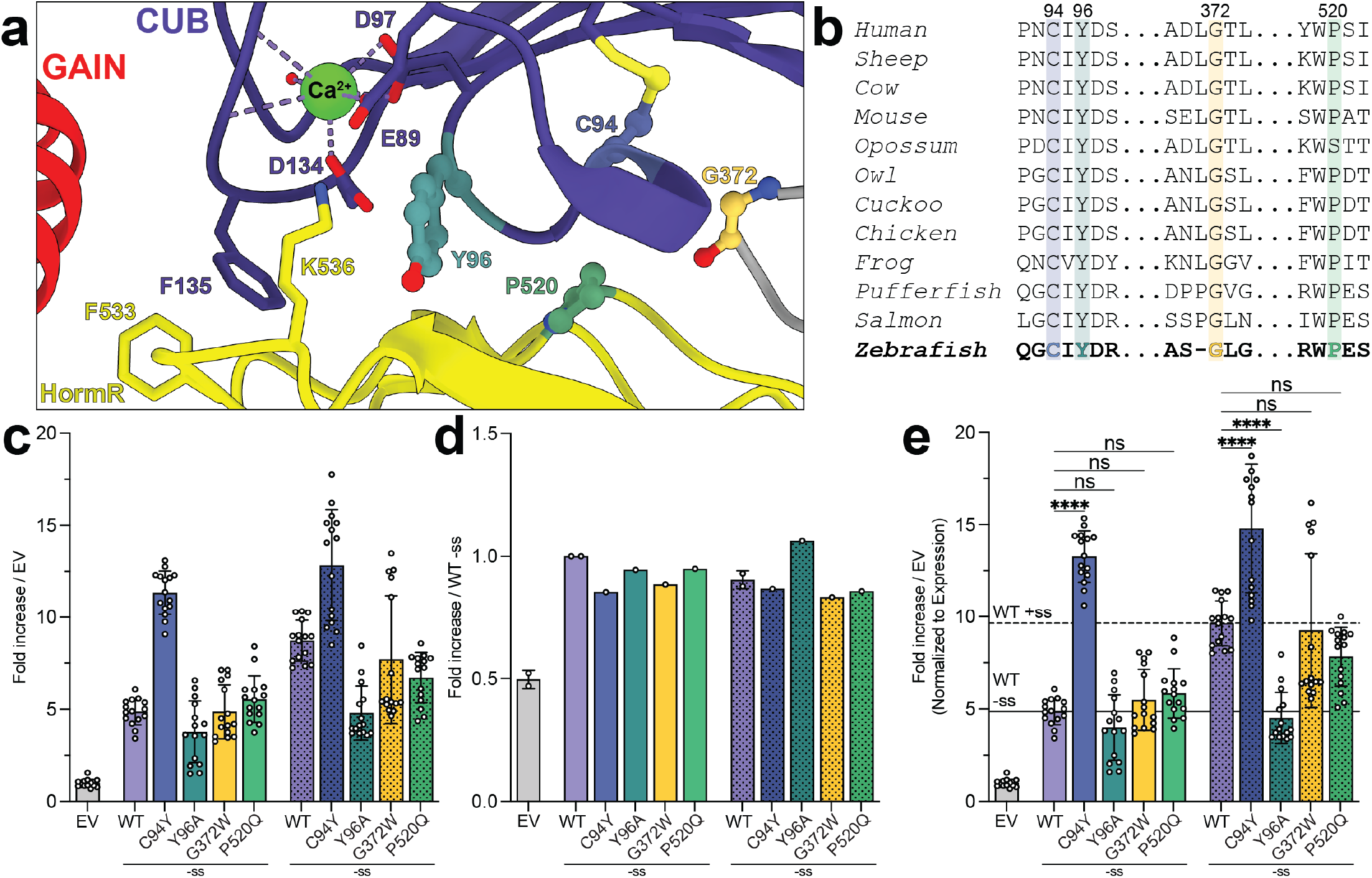
Point mutants sourced from cancer patients modulate ADGRG6 cAMP signaling. **(a):** Close–up of the CUB:HormR interface from the crystal structure with key residues involved in the interaction shown as sticks. Residues mutated in cancer patients are represented as differently colored spheres (C94 light purple,Y96 teal, P520 green, and G372 gold). The bound Ca^2+^ ion is shown as a green sphere with coordination shown as dotted lines. **(b):** Sequence conservation of residues in the CUB:HormR interface is shown using ADGRG6 orthologs. Cancer-associated residues are colored as in (a). **(c):** Basal cAMP signaling of ADGRG6 constructs shown as fold change over empty vector (EV). Mutants are grouped by absence (–ss) or presence (+ss) of the ECR splice insert. N=3 independent experiments were performed with five technical replicates each. **(d):** Relative cell surface expression levels for ADGRG6 constructs shown as fold change over WT –ss. See Fig. S6A, B for raw flow cytometry data. N=1 independent experiments were performed. **(e):** Basal cAMP signaling represented as fold change over EV normalized to cell–surface expression with cancer mutants colored as in (a). Horizontal lines show the mean normalized signaling levels for WT constructs. **** – p<0.0001, ns – not significant; one–way ANOVA test with Tukey’s correction for multiple comparisons. Each condition was compared to every other condition, but not all comparisons are shown for clarity. The following comparison (WT –ss vs. WT +ss) results in ****, p<0.0001 and (C94Y –ss vs. C94Y +ss) results in ns but these are not shown for clarity. N=3 independent experiments were performed with five replicates each. Bar graphs represent mean values with the standard deviation being shown for error.

Similar to seen previously (Fig. 6E) (14), the ADGRG6 –ss isoform signals 5–fold over empty vector (EV) and ADGRG6 +ss signals at a 10–fold level over EV, a significantly higher extent than –ss (Fig. 6E). ADGRG6 C94Y was found super active (∼ 14–fold), with both isoforms signaling higher than even the WT +ss receptor. Furthermore, there is no significant difference between C94Y –ss and +ss signaling levels, in contrast to the isoform-dependence of WT ADGRG6 signaling. The Y96A –ss mutant signaled at a level comparable to WT –ss (Fig. 6E). However, Y96A +ss signaling was 5–fold over EV, similar to WT –ss. This decrease in signaling suggests that the event in WT +ss which leads to elevation of cAMP is not possible in Y96A. G372W and P520Q ADGRG6 were not significantly different from WT (Fig. 6E). Taken together with our computational studies, these results suggest that the dissociation of CUB from the HormR domain may activate ADGRG6, but only if residues in the Ca^2+^ binding surface are unaffected. In our system, WT and C94Y ADGRG6 were similarly activated by the TA peptide added in trans (Fig. S6C, D). Also, WT and C94Y ADGRG6 were both insensitive to addition of reported ligands (Fig. S6I, J), including PrP^c^ (36) and CIV (35).

## Discussion

Despite their essential functions in development and their role in disease, aGPCRs remain an understudied subfamily of cell–surface receptors with enormous therapeutic potential (5, 6). Understanding the molecular basis of aGPCR function is an important step towards therapeutic development. Within the aGPCR family, ADGRG6 has gained attention due to its roles in PNS myelination, angiogenesis, heart and skeletal development, as well as its association with developmental diseases and cancers (22–26). Overall, our results support an ECR–dependent mechanism for ADGRG6 regulation that centers around the far N–terminal CUB domain. Our results advance previous models of ADGRG6 regulation and elucidate a CUB-centered mechanism that may guide future therapeutic development.

### The far N-terminal CUB domain of ADGRG6 as a key regulator of signaling

Our data show that the CUB domain of ADGRG6 regulates cAMP signaling. Exposure of the CUB domain to solvent increases receptor signaling, and the residues in the Ca^2+^ binding surface need to be intact for the regulatory role of the CUB domain (Fig. 7). We demonstrate that the ΔCUB splice variant produces receptors which have decreased cAMP signaling. In this case, CUB cannot bind to any ligand, intramolecular or otherwise (Fig 7A). For WT ADGRG6 –ss, the ECR samples closed conformations with the CUB domain bound to the HormR domain (Fig. 7B), and a moderate level of signaling is observed. We speculate that CUB binding its intramolecular HormR domain stabilizes an ECR conformation which drives this moderate level of signaling. WT +ss and C94Y favor exposure of the Ca^2+^ binding site to solvent but leave the Ca^2+^ binding surface unaffected (Fig 7C), and we observe these versions of the protein in a higher signaling state than WT –ss. We propose that the open-like ECR conformations, such as those observed in our MD simulations of C94Y, transmit a conformational change to the 7TM that drives this higher signaling.

**Figure 7.**
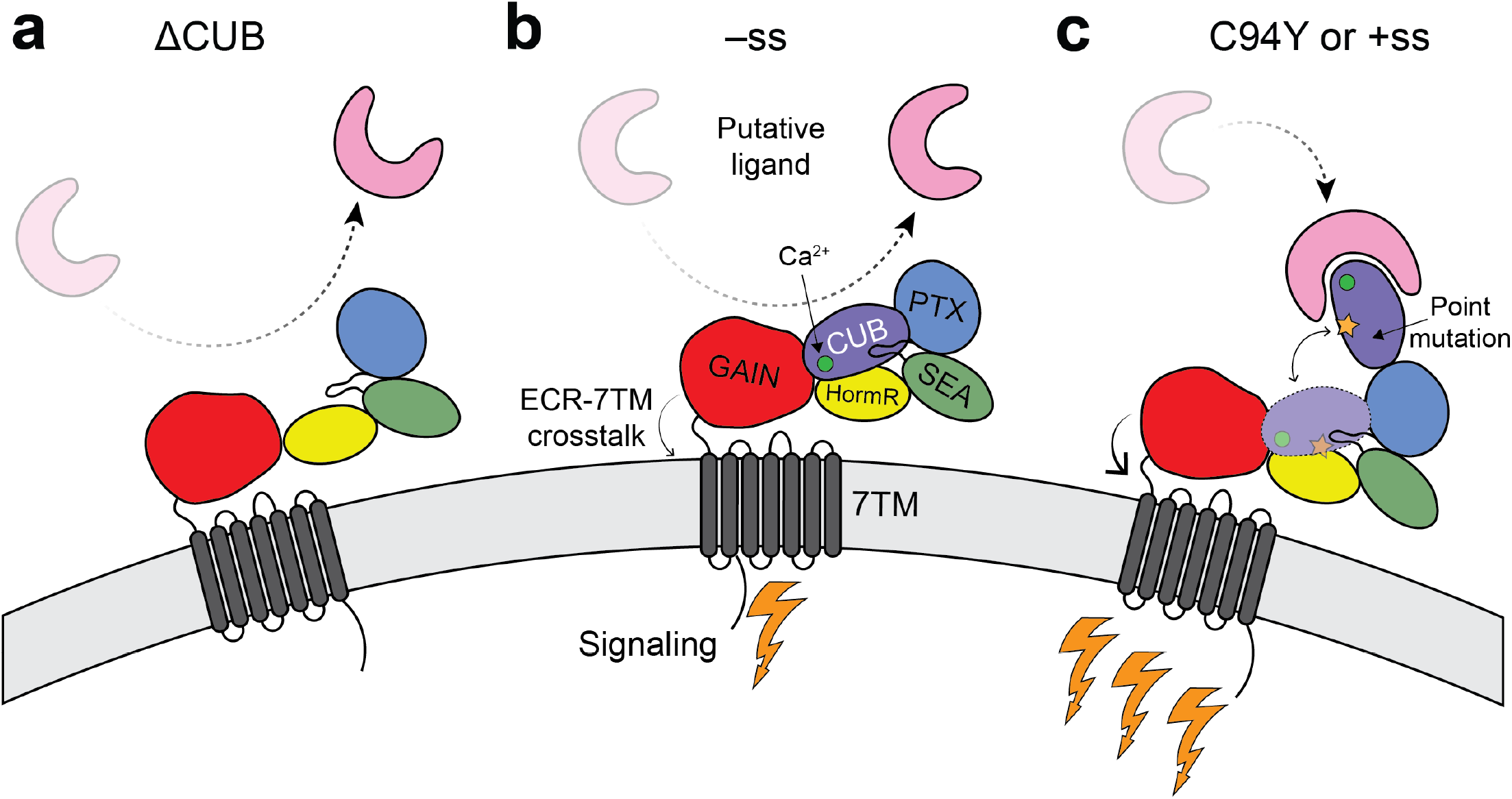
Model for regulation of ADGRG6 signaling by its CUB domain. ADGRG6 is shown as a cartoon model with the putative ligand shown as a pink U–like shape and signaling shown as golden lightning bolts. A bound calcium ion is depicted as a green circle inside the CUB domain. The point mutation in the CUB domain is shown as a golden star. More lightning bolts indicate a higher level of signaling. The TA cleavage site is shown using scissors inside the GAIN domain. **(a):** The ΔCUB receptor is unable to either bind the intramolecular HormR site or a putative extracellular ligand and has significantly decreased cAMP signaling (no lightning bolts). **(b):** For the ADGRG6 –ss receptor, the CUB/HormR interface remains intact. The –ss receptor signals modestly (one lightning bolt) and the CUB domain is unable to bind a putative extracellular ligand. **(c):** Mutants which break the CUB/HormR interface but not the canonical CUB ligand binding site displace the CUB domain from HormR. The receptor then accesses a higher signaling state (three lightning bolts) upon conformational change, which includes potential ligand binding events.

Finally, the Y96A mutation behaves similarly to the previously characterized D134A/F135A Ca^2+^ binding surface mutation (14) in our signaling experiments, as neither of these mutants access the higher signaling levels available to WT +ss or C94Y. Both Y96A and D134/F135A are part of the ADGRG6 Ca^2+^ binding surface and are key to ligand binding in other CUB domain-ligand complexes (32–34). These results ascribe further importance to the highly conserved Ca^2+^ binding surface on CUB as a regulator of ECR-dependent signaling.

### The aGPCR ECR as an allosteric sensor of ligand binding

Our work on ADGRG6 joins a growing body of work which altogether suggests that aGPCRs use N-terminal portions of their ECRs to sense ligand binding or force; these events can then be transmitted by the ECR to the 7TM to alter signaling. ADGRL3 uses its N-terminal lectin and olfactomedin domains to interact with TEN2 and FLRT (9, 38–40) to instruct synapse specificity, and anti-lectin synthetic antibodies (sABs) can modulate signaling (10). Furthermore, for ADGRL3, anti-GAIN sABs which alter the GAIN/7TM relative orientation can affect receptor signaling (19), supporting an allosteric connection between ECR and 7TM in aGPCRs. ADGRG1 directs myelination through the ligand of its N-terminal PLL domain, transglutaminase-2 (41, 42), and anti-PLL monobodies can alter signaling (12). ADGRC1/CELSR1 uses its N-terminal cadherin repeats to mediate homophilic adhesion in trans (18, 43), and may use a compact module of several domains to transmit this signal to the 7TM (18). The N-terminal PTX domain of ADGRG4 is proposed to dimerize and sense shear force which may result in ADGRG4 activation (44).

In regards to ADGRG6, this receptor interacts with its putative ligand Collagen IV through the CUB–PTX region of its ECR leading to elevation of cAMP signaling levels (35). Furthermore, two alternatively spliced versions of ADGRG6 both modulate the CUB Ca^2+^ binding surface, providing support for its importance as an ADGRG6 regulatory hotspot. The ΔCUB splice variant removes the entire CUB domain (26), preventing the Ca^2+^ binding surface from binding to any ligand, and the +ss splice insert leads to increased solvent exposure of the Ca^2+^ binding site (14), disrupting the closed conformation, and allowing it to interact with ligands other than HormR.

Since several ADGRG6 ligands are reported in the literature (29, 35, 36), it is possible that the Ca^2+^ binding surface on CUB interacts with these ligands or others to regulate receptor function. Additionally, it is possible that conformational changes in the ECR upon disruption of the closed conformation yield the higher signaling activity observed for WT +ss or C94Y. Future structural and mechanistic work will clarify how the CUB Ca^2+^ binding surface modulates receptor function, and perhaps generate tool molecules such as antibodies to bind to the CUB domain and modulate ADGRG6 signaling. Such reagents may have the ability to serve as activators or inhibitors of ADGRG6 signaling and will lay the groundwork for therapeutic development.

## Acknowledgements

This work was supported by a National Institutes of Health Grant R35GM148412 (to DA) and a National Institutes of Health Grant F32GM142266 and K99GM157487 (to SJB). GM/CA@APS has been funded by the National Cancer Institute (ACB-12002) and the National Institute of General Medical Sciences (AGM-12006, P30GM138396). This research used resources of the Advanced Photon Source, a U.S. Department of Energy (DOE) Office of Science User Facility operated for the DOE Office of Science by Argonne National Laboratory under Contract No. DE-AC02-06CH11357. The Eiger2 16M detector at GM/CA-XSD was funded by NIH grant S10OD034267. We thank the CCP4 School in Macromolecular Crystallography for assistance in determination of the crystal structure. This work was completed in part with resources provided by the University of Chicago Research Computing Center. The content is solely the responsibility of the authors and does not necessarily reflect the official views of the National Institute of General Medical Sciences or the National Institutes of Health.

## Author contributions

Conceptualization: ED, SJB, DA, RV, KC. Methodology: ED, SJB, DA. Investigation: ED, SJB, KL, SPK, TS. Visualization: ED, SJB. Supervision: DA. Writing-original draft: ED. Writing-review & editing: ED, SJB, DA.

## Materials and Methods

### Molecular dynamics simulations

Software used for this work including GROMACS (45) was compiled and maintained by SBGrid (46). Simulations were initialized from PDB 6V55 (14). Atoms missing from the crystal structure were added with Swiss PDB Viewer. N–linked glycosylation and crystal structure waters were removed from the protein. Using GROMACS, the structure was placed in a 4216.9 nm^3^ box with periodic boundary conditions such that all protein atoms are at least 13 Å from the boundaries, resulting in a 160 × 160 × 160 Å cube (Fig. S2A). The box was solvated with the SPC water model using `gmx solvate`and Na^+^ and Cl^−^ counterions were added using `gmx genion`. The resulting model of ADGRG6 was minimized with the leapfrog integrator in step sizes of 0.1 Å until the max force was smaller than 1000 kJ mol^−1^ nm^−1^. The system was then subjected to a two–phase equilibration in the NVT and NPT ensembles with protein atoms under harmonic position restraints. A 100 ps NVT equilibration was carried out with step size 2 fs at 300 K with the Berendsen thermostat using a 0.1 ps coupling constant, and solvent velocities were assigned from the Boltzmann distribution. NPT equilibration was also carried out with step size 2 fs for 100 ps, and at 1 atm with Parrinello–Rahman pressure coupling (2.0 ps coupling constant) using velocities from the NVT ensemble. Simulations on the unrestrained protein were run in the NPT ensemble for 1 µs and a 2 fs step size. Long–range electrostatics were computed with PME with a 1.6 Å Fourier spacing and 10 Å cutoff. All simulations were run with the AMBER99SB–ILDN force field. Replicate simulations were each separately solvated, minimized, and equilibrated.

Simulated Systems
| Genotype | Splice isoform | Simulation Length | Replicates |
| --- | --- | --- | --- |
| WT | –ss | 1 μs | 5 |
| C94Y | –ss | 1 μs | 3 |

### Simulated Systems

### PC analysis and free energy landscape analysis

Trajectories from the simulations were extracted for protein atoms and output saved for every ten frames. PCs of the Cαs were calculated by concatenating replicate simulation trajectories for each condition using `gmx trjcat`and diagonalizing the covariate matrix with `gmx covar`. Trajectories were projected onto PCs using `gmx anaeig`and were plotted and visualized with a homemade python script. The fractions of variance *f* captured by PCs were checked to verify that the first two principal components captured most of the variance and verify the quality of the dimensionality reduction by computing:

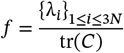

where {λi}_1≤i≤3N_ is the set of 3*N* eigenvalues for *N* α–carbons, and tr(*C*) is the trace of the covariate matrix. For visualization of trajectories over time on PC space, a Gaussian filter with σ = 50 was applied to PC1 and PC2 components of each simulation’s trajectory. Free energy landscapes were obtained by binning the unfiltered projected simulations into 50 × 50 bins and computing the Gibbs free energy (*G*_*i,j*_) for each bin:

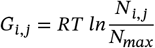

where *N* _*i,j*_is the number of frames in the *i*^th^ bin of PC1 and *j*^th^ bin of PC2 and *N*_*max*_ is the number of frames in the most occupied bin. Hierarchical clustering of free energy landscape wells was performed by scipy.cluster in a homemade python script using the Ward clustering method to minimize variance in in the *l*^2^ norm. Cluster centroids *c*_*i*_ were compsssssuted by averaging the PC1 and PC2 coordinates of frames within each cluster:

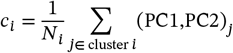

where *N _i_*is the number of frames in the *i*^th^ cluster. We chose the frame from the simulations whose PC coordinates are closest by *l*^2^ norm and constructed the 3D structure as a linear combination of PC1 and PC2 using gmx anaeig. For entropy calculations, we used Schlitter’s formula due to the size of the system and emphasis on higher inertia eigenvectors to approximate an upper bound for the protein’s entropy (*S*_system_) as follows:

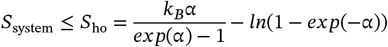

where:

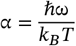

*S*_ho_ is the entropy of the harmonic oscillator, and ω is the frequency of the harmonic oscillator related to the variance of the system. Entropy calculations were performed with gmx anaeig –entropy. Cumulative entropy plots were calculated at each time *t* from the covariate matrix for the portion of the simulation [0, *t*]. Subspace overlap between two different simulations with associated sets of eigenvectors *v*_1_, *v*_2_, …, *v*_3*N*_ and ***w***_1_, ***w***_2_, …, ***w***_3*N*_ is found by:

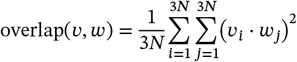

producing overlap(***v, w***) ∈ [0,1]. Computation of overlap was carried out with `gmx anaeig –over`.

### Cloning

Wild–type and mutant *D. rerio* (zebrafish) ADGRG6 (14) (numbered in this manuscript according to Uniprot ID: C6KFA3) constructs were cloned into pCMV5 by annealing three fragments with the Gibson assembly method. All constructs contain residues 41–1185 and include N–terminal FLAG–tags for measuring cell–surface expression levels. pCMV5 backbone was amplified using the following primers: F: 5’ – CGTCCAC-CAGAAAGGAGCAGTGG – 3’, R: 5’– GGGAACCGGAGCTGAATGAA-GCC – 3’. The first ADGRG6 –ss Y96A fragment was mutated and amplified from full–length ADGRG6 DNA using: F: 5’ – AGCACAAGGCTG-CATGCGTGACCGCGTTGTCGTC – 3’, R: 5’ – CCAC-TGCTCCTTTCTGGTGGACG – 3’. The second Y96A fragment was mutated and amplified from full–length ADGRG6 DNA using F: 5’ – GGCTTCATTCAGCTCCGGTTCCC – 3’ and R: 5’ – GACGACAAC-GCGGTCAGCGATGCAGCCTTGTGCT – 3’. In a similar scheme to Y96A, the following fragment: primer combinations were utilized to amplify other point mutants (5’ – 3’). C94Y, 1 GAGGAAGCACAAGGC-TACATCTATGACCGCG, CCACTGCTCCTTTCTGGTGGACG. C94Y, 2 GGCTTCATTCAGCTCCGGTTCCC, CGCGGTCATAGATGTAGCCTT-GTGCTTCCTCG. P520Q, 1 CCACTATAGATGGCAAGAGAG-CAGACCCACAG, CCACTGCTCCTTTCTGGTGGACG. P520Q, 2 GCTTCATTCAGCTCCGGTTCCC, GACTGTGGGTCTGCTCTCTT-GCCATCTATAGTGG. G372W, 1 GCAGGTTGTGCTTCTTGGCTT-GGCTGCCCAG CCACTGCTCCTTTCTGGTGGACG. G372W, 2 GGCTTCATTCAGCTCCGGTTCCC, CTGGGCAGCCAAGCCAAGAA-GCACAACCTGC. The identity of each construct was confirmed with full–plasmid sequencing. Δ CUB constructs which contain ADGRG6 residues 149–1185 were generated using an inverse PCR scheme. WT ADGRG6 plasmids were subjected to inverse PCR using the following primers: 5’ AAGCTTGTCGTCATCGTCTT 3’ and 5’ GCAG-TGACTCTGAGGAACCA 3’. The resulting linear product contained the entire source plasmid except for the CUB domain. This linear product was circularized using Gibson assembly. The identity of constructs were confirmed using Sanger sequencing.

### cAMP signaling assay

HEK293 cells (ATCC CRL–1573) were seeded in 6–well plates with Dulbecco’s Modified Eagle Medium (DMEM; Gibco, 11965092) supplemented with 10 % FBS (Sigma–Aldrich, F0926). At 60–70 % confluency, the cells were co–transfected with 0.35 µg Gpr126 DNA, 0.35 µg GloSensor reporter plasmid (Promega, E2301), and 2.8 µL transfection reagent Fugene 6 (Promega, PRE2693). After a 24–h incubation, the transfected cells were detached and seeded (50,000 cells per well) in a white 96–well assay plate. Following another 24–h incubation, the DMEM was replaced with 100 µL Opti–MEM (Gibco, 31985079) and incubated for 30 min. To each well was then added 1 µL GloSensor substrate and 11 µL FBS, and 0.5 µg/mL Collagen IV (Sigma Aldrich CC076, dissolved in 1x PBS) or 4 µM of the flexible tail of PrP^c^ (residues 23-34 (36) were synthesized by GenScript and dissolved in 1x PBS) or the equivalent volume of PBS as a control. Basal–level luminescence measurements were taken after 30 min to allow for equilibration. For TA peptide addition, after basal signaling had stabilized, 100 µM p14 TA was added (GenScript; in 100 % DMSO) or the equivalent volume of DMSO as a control.

### Flow cytometry

HEK293T cells were transfected in a 6–well plate with 1 µg ADGRG6 DNA and 3 µL transfection reagent Fugene 6 (Promega, PRE2693) at 60–70% confluency and incubated for 48 h. The cells were detached with citric saline and washed with phosphate–buffered saline (PBS). The cells were then washed twice with PBS supplemented with 0.1% BSA (Sigma–Aldrich, A3803). The cells were incubated with mouse anti–FLAG primary antibody (1:1000 dilution in PBS + 0.1% BSA; Sigma–Aldrich, F3165) at room temperature for 30 min and washed three times with PBS + 0.1% BSA. The cells were then incubated with donkey anti–mouse Alexa Fluor 488 secondary antibody (1:500 dilution in PBS + 0.1% BSA; Invitrogen) at room temperature for 30 min and washed three times with PBS + 0.1% BSA. Stained cells were resuspended in PBS + 0.1% BSA and at least 5000 data points were collected with a BD Accuri C6 flow cytometer. The resulting data were analyzed using FloJo v10, single HEK293T cells were gated, and median fluorescence intensity in the 488 channel was used.

### Statistical Analysis

All statistical analysis for the cAMP signaling assays was performed using GraphPad Prism Version 10. All experiments were performed in at least N=3 independent experiments each with at least three replicates. For statistical comparisons, the one–way ANOVA with Tukey’s correction for multiple comparisons test was used. Each condition was statistically compared with every other condition, but only some comparisons are shown for clarity.

