## Supplemental Information for "The far extracellular CUB domain of the adhesion GPCR ADGRG6/GPR126 is a key regulator of receptor signaling"

**This PDF file includes:**

Supporting information
Figures S1 to S6
Table S1
Legends for Movies S1 to S6
SI References

**Other supporting materials for this manuscript include the following:**

Movies S1 to S6

### Supporting Information Text

#### Supporting Methods

##### Cell culture, protein purification, and protein crystallography

High Five insect cells (Trichoplusia ni, female, ovarian. ThermoFisher, B85502) were used for production of recombinant proteins. Cells were cultured using Insect-Xpress medium (Lonza, 04351Q) with 10 μg/mL gentamicin at 27 °C. Sf9 cells (Thermo Fisher, 12659017) were transfected with ADGRG6 +ss DNA and commercial baculovirus DNA (Expression Systems, 91- 002) using Cellfectin II (Thermo Fisher, 10362100). High-titer recombinant baculovirus was obtained using Sf9 cells grown in SF900-III medium with 10% (v/v) FBS (Sigma-Aldrich, F0926). High Five cells (Thermo Fisher, B85502) were infected with baculovirus at $2.0 \times 10^6$ cells/mL and incubated at 27 °C with 120 rpm shaking for 72 h. ADGRG6 +ss (residues S38-A839) was cloned with an 8x polyhistidine tag at the C-terminus. Media was harvested from cell cultures 72 hours after infection. The media was centrifuged at room temperature at 900 x g for 15 min. The supernatant was harvested and transferred to a beaker with stirring at room temperature and the following were added: 50 mM Tris pH 8.0, 5 mM $CaCl_2$ and 1 mM $NiCl_2$ (final concentrations listed). After 30 min, the solution was centrifuged for 30 min at 8000 x g. The clarified supernatant was then incubated with nickel-nitriloacetic (Ni-NTA) resin (QIAGEN 30250) with stirring at room temperature for 3 hours. A Büchner funnel was used to collect resin and wash using a buffer composed of 10 mM HEPES pH 7.2, 150 mM NaCl (HBS) and 20 mM imidazole, and then the washed resin was transferred to a poly-prep chromatography column (Bio-Rad). The protein was eluted using HBS buffer containing 200 mM imidazole. Fractions containing desired protein were pooled and concentrated using a 50 kDa centrifugal concentrator (Amicon UFC805024) and loaded on gel-filtration chromatography using a Superdex 200 10/300 column (GE Healthcare) in HBS. Purified ADGRG6 +ss was concentrated to 20 mg/mL using a 50 kDa cutoff centrifugal concentrator system (Amicon UFC805024) and applied to sitting drop crystal trays (Fig. S2A) in a drop volume of 0.2 μL against a well volume of 50 μL. Crystals were grown in a well solution consisting of 0.1 M HEPES pH 7.0 with 1.1 M Na Citrate (Fig. S2B). Crystals grew at room temperature. Crystals were looped from their mother liquor and transferred to a cryoprotection solution of original mother liquor supplemented with 20 % Ethylene Glycol and then immediately frozen in liquid nitrogen. Frozen crystals were shipped on liquid nitrogen to the GM/CA beamline at the Advanced Photon Source at Argonne National Labs for remote data collection at 110 K (Fig. S2C, D). Final resolution was determined using shells which contain $CC_{1/2} > 0.2$ using analysis done by aimless (1). Detailed data collection and processing statistics are available in Table 1.

##### Data processing and structure determination

Data collected at GM/CA were reduced, indexed, integrated, and scaled using HKL2000. All structural solution and refinement steps were carried out using the PHENIX suite of software for macromolecular crystallography (2). Automated solutions suggested by the beamline were in $C222_1$ and this space group yields reasonable molecular replacement solutions in PHASER using the GAIN-HormR-SEA tridomain structure from PDB 6V55 (TFZ 13.3 – 1 copy per ASU). However, upon manual inspection of the crystal packing, obvious clashes are present which make these unlikely solutions. Regardless, phenix.refine was attempted and yielded $R_{free}$=0.43. Using this partial solution the CUB and PTX domains could not be placed, despite ample room in the unit cell. We also tried lower symmetry space groups and the best solution we obtained was in space group $C_2$ with 2 molecules of GAIN-HormR-SEA per asymmetric unit (TFZ 23.1; $R_{free}$=0.39). (RMSD= 0.65 Å from the -ss ADGRG6 structure) and several other indications that the solution is correct (sensical crystal contacts, reasonable $R_{free}$ for the estimated resolution, difference density for several N-linked glycosylation sites (Fig. S2J). Using this improved solution, we once again tried to place the CUB and PTX domains using molecular replacement but could not, despite available space in the unit cell.

Since we do not observe any density for the CUB/PTX domains (or anything outside of GAIN-HormR-SEA), this is consistent with the idea that CUB/PTX may be present, but flexible relative to SEA/HormR/GAIN. Matthews probability analysis suggests that given the best obtained solution (space group $C_2$ with 2 molecules of GAIN-HormR-SEA per asymmetric unit), it is ~10 times more likely that the entire ECR sequence (90 kDa) is present in the unit cell vs only the resolved sequence (50 kDa) being present (Fig. S2 E vs. F). There is enough space in the crystal lattice to accommodate the CUB/PTX domains (Fig. S2). Thus, we suspect that the most likely scenario is that the CUB/PTX domains are indeed present in this structure, but flexible and unresolved, relative to the resolved GAIN-HormR-SEA domains.

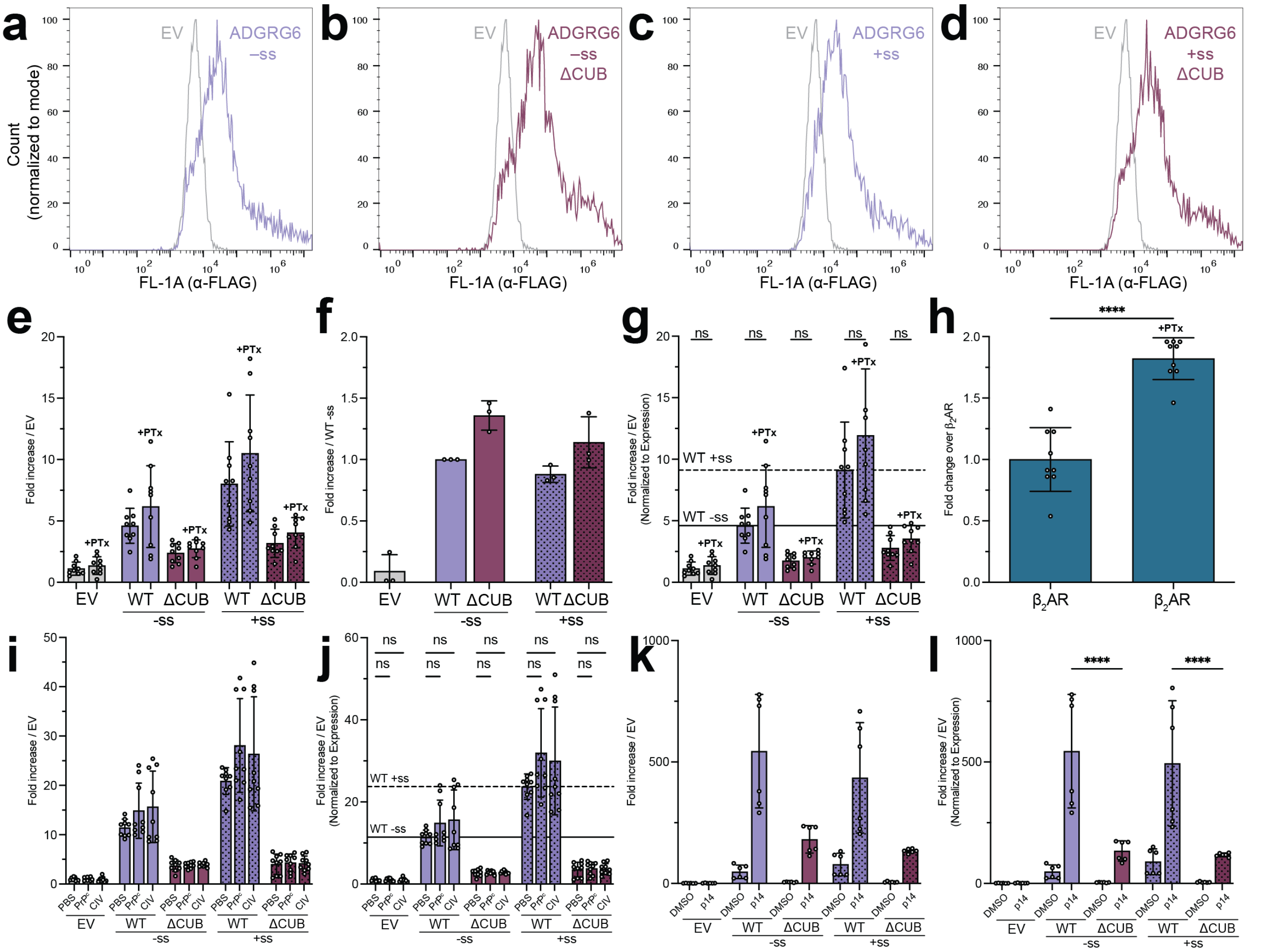


**Fig. S1. Expression levels of ADGRG6 ΔCUB constructs and investigation of ligand-stimulated ADGRG6 activity. (a)-(d):** Representative flow cytometry data showing expression levels of the N-terminally FLAG-tagged constructs used for cAMP signaling assays shown in Figure 1. These are measured using AlexaFluor-488 labeled α-FLAG antibodies. Each plot is overlaid with EV from the same experiment to highlight background labeling of cells. The cell count in the Y axis is normalized to the mode of each dataset. Median FL-1A (α-FLAG) values are used to quantify the cell-surface expression of each construct. **(e):** Basal cAMP signaling of ADGRG6 constructs shown as fold change over empty vector (EV) in response to 500 ng/mL pertussis toxin (PTx) addition. Mutants are grouped by absence (–ss) or presence (+ss) of the ECR splice insert. N=3 independent experiments were performed with three technical replicates each. **(f):** Relative cell surface expression levels for ADGRG6 constructs shown as fold change over WT –ss. N=3 independent experiments were performed. **(g):** Basal cAMP signaling from (e) represented as fold change over EV normalized to cell-surface expression. Horizontal lines show the mean normalized signaling levels for WT constructs. ns – not significant; one–way ANOVA test with Tukey's correction for multiple comparisons. Each condition was compared to every other condition, but not all comparisons are shown for clarity. N=3 independent experiments were performed with three replicates each. Bar graphs represent mean values with the standard deviation being shown for error. **(h):** The $\beta_2$ Adrenergic receptor ($\beta_2$AR), a well characterized receptor which couples to both G$\alpha_s$ and G$\alpha_i$, exhibits increased cAMP levels upon PTx addition, serving as a positive control for PTx function. **** – p<0.0001; one–way ANOVA test with Tukey's correction for multiple comparisons. **(i):** Basal cAMP signaling of ADGRG6 constructs shown as fold change over empty vector (EV) in response to addition of Phosphate Buffered Saline (PBS) as a loading control, the flexible tail of the cellular prion protein (PrP$^c$), or Collagen IV (CIV). Constructs are grouped by absence (–ss) or presence (+ss) of the ECR splice insert. N=3 independent experiments were performed with three technical replicates each. **(j):** Basal cAMP signaling from (i) represented as fold change over EV normalized to cell-surface expression. Horizontal lines show the mean normalized signaling levels for WT constructs. ns – not significant; one–way ANOVA test with Tukey's correction for multiple comparisons. Each condition was compared to every other condition, but not all comparisons are shown for clarity. N=3 independent experiments were performed with three replicates each. Bar graphs represent mean values with the standard deviation being shown for error. **(k):** Basal cAMP signaling of ADGRG6 constructs shown as fold change over empty vector (EV) in response to addition of Dimethyl Sulfoxide (DMSO) as a loading control or the 14-residue tethered agonist peptide of ADRGR6 (p14). Constructs are grouped by absence (–ss) or presence (+ss) of the ECR splice insert. N=2 independent experiments were performed with three technical replicates each. **(l):** Basal cAMP signaling from (i) represented as fold change over EV normalized to cell-surface expression. **** – p<0.0001; ns – not significant; one–way ANOVA test with Tukey's correction for multiple comparisons. Each condition was compared to every other condition, but not all comparisons are shown for clarity. N=2 independent experiments were performed with three replicates each. Bar graphs represent mean values with the standard deviation being shown for error.

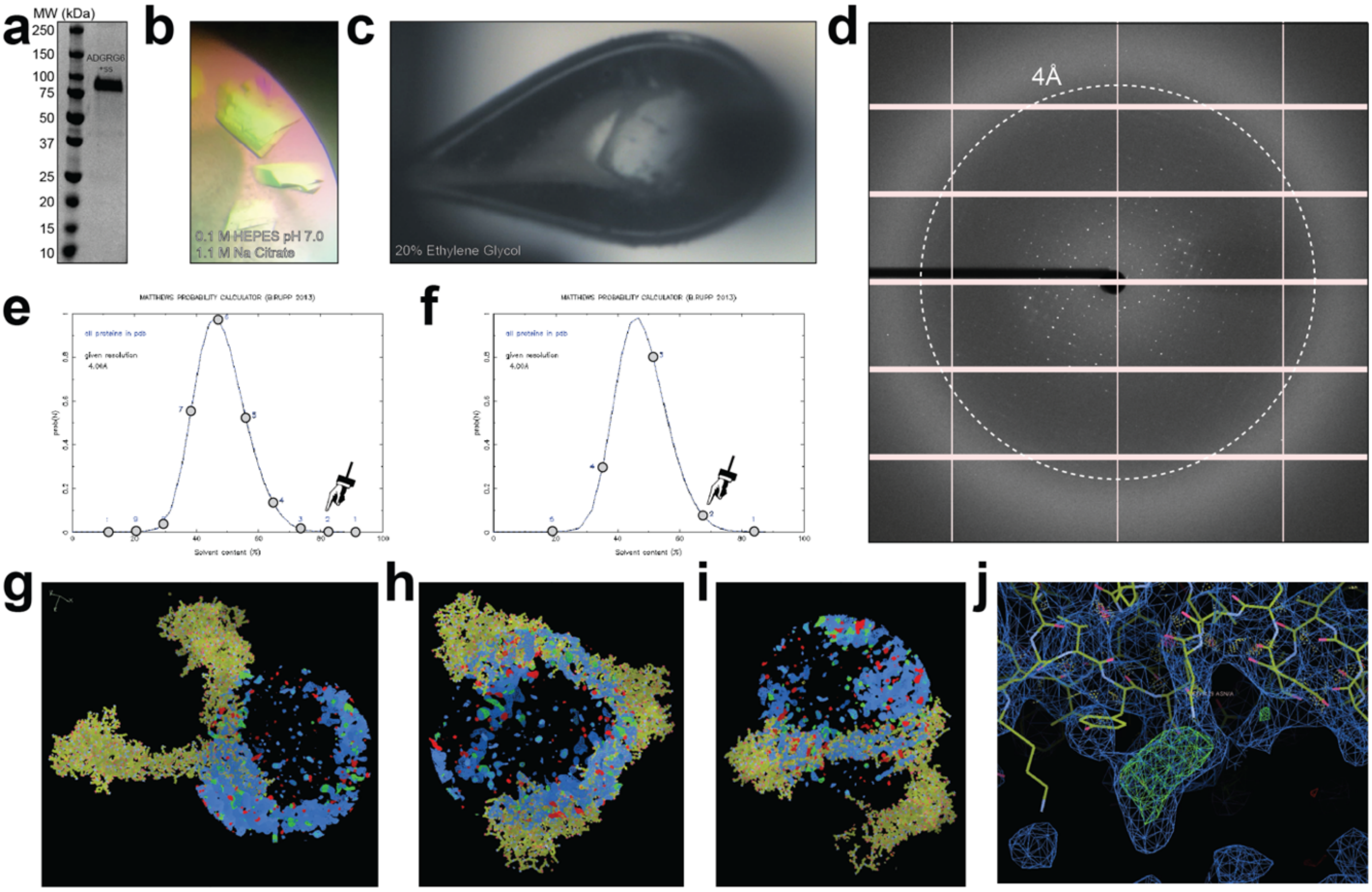


**Fig. S2. The crystal structure of the ADGRG6 +ss ECR suggests that the CUB and PTX domains are dynamic relative to the SEA/HormR/GAIN domains. See methods and table 1 for more information. (a):** Reducing SDS-PAGE of the purified ADGRG6 +ss ECR. **(b):** Protein crystals of the ADGRG6 +ss ECR were grown using hanging drop vapor diffusion. **(c):** ADGRG6 crystals were cryoprotected using the mother liquor plus 20 % Ethylene Glycol and sent to the GM/CA beamline 23-ID-D. **(d):** Representative diffraction image shows visible spots extending to 4Å. **(e, f):** Calculation of Matthews coefficient probabilities (the best obtained solution is C2 with two copies per ASU) suggest that the lattice is ~ 10 times less likely to form if the sample were cleaved at the ordered portion vs. having the two unobserved domains present but disordered. **(g, h, i):** Different views of the crystal lattice shows that there is no observed density for the CUB or PTX domains in the lattice. **(j):** Factors such as difference density at N-linked glycosylation sites suggest that this is the correct solution.

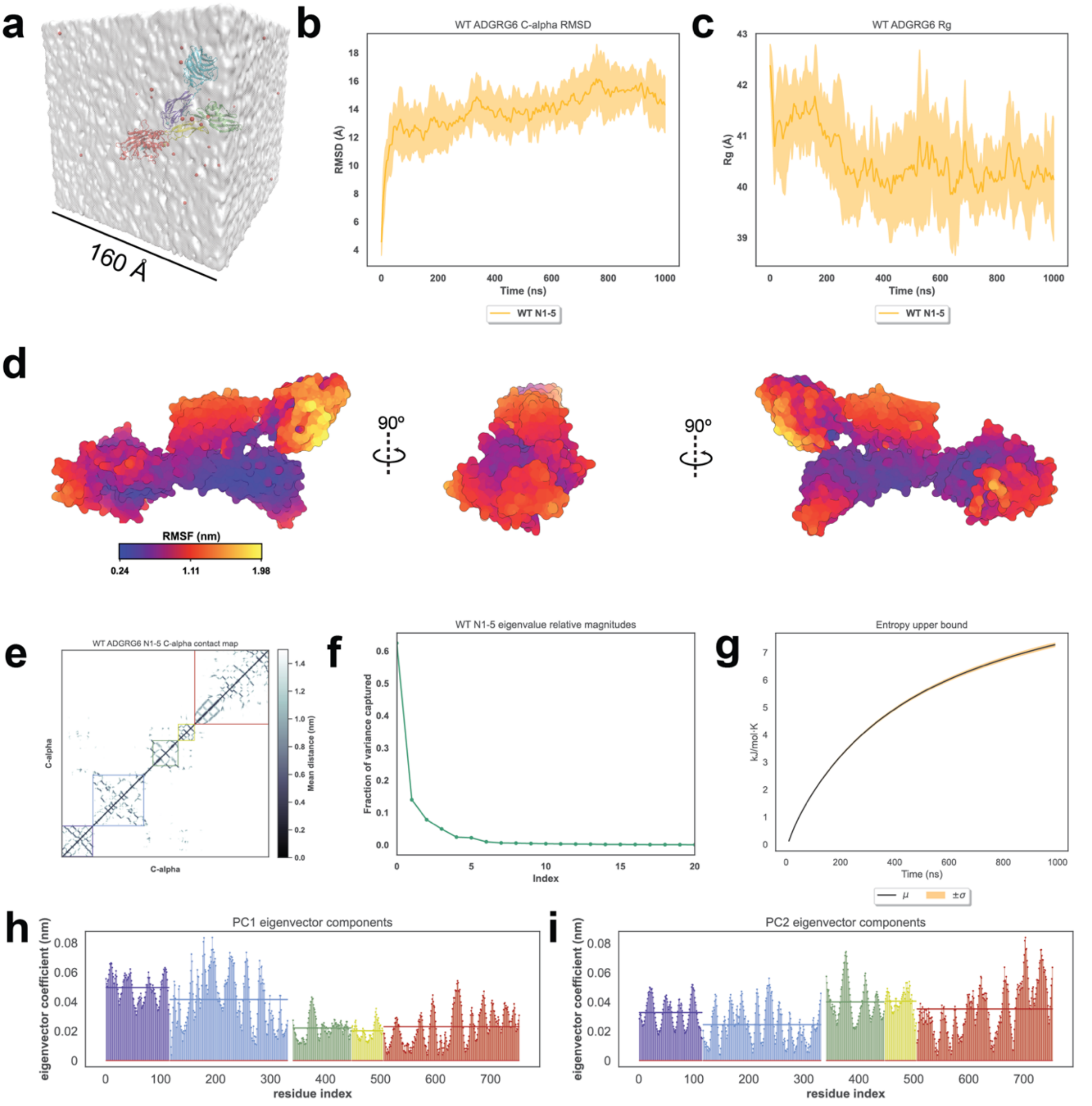


**Fig. S3. Dynamics of the WT zfADGRG6 –ss ECR throughout five one–microsecond MD simulations. (a):** System simulated for each MD experiment, with cubic water box displayed and sodium counterions represented as red spheres. **(b):** Average root-mean-square deviation (RMSD, dark line) and standard deviation (shaded region) of the Cα atoms from the crystal structure computed across the five simulations. **(c):** Radius of gyration ($R_g$) mean (dark line) and standard deviation (shaded region) across all five simulations. **(d):** Root-mean-square fluctuation (RMSF) from the average position for each residue mapped onto a surface representation of the average zfADGRG6 –ss ECR simulation structure shown from three different views. **(e):** Mean distance matrix between each alpha carbon computed across all five simulations. Colored boxes denote domain boundaries, and data outside of colored boxes represent interdomain contacts. **(f):** Scree plot showing the variance of the concatenated WT simulations captured by each PC. Only the first 20 of 2259 PCs are shown for clarity. **(g):** Cumulative entropy upper bound across all five simulations computed by Schlitter's formula. Black line denotes mean and red boundaries denote the standard deviation. **(h):** Magnitude of the Cartesian components for each Cα in PC1 colored by domain to represent the conformational variance of each domain. Colored horizontal lines denote mean magnitudes for each domain. **(i):** Magnitude of the Cartesian components for each Cα in PC2 represented as in (h).

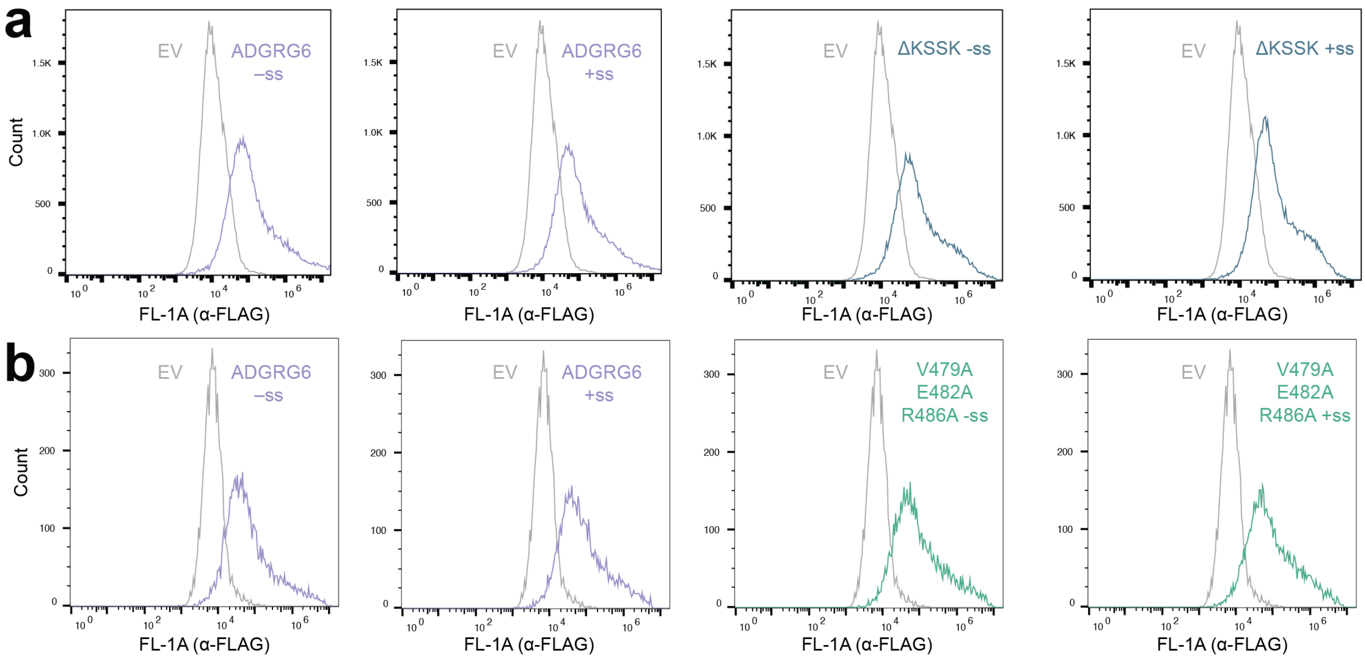


**Fig. S4. Expression levels for ADGRG6 PTX/SEA interface mutants.** Flow cytometry data showing expression levels of the N–terminally FLAG–tagged constructs used for cAMP signaling assays in Figure 5. These are measured using AlexaFluor–488 labeled α–FLAG antibodies. Each plot is overlaid with EV from the same experiment to show background labeling of cells. Median FL-1A (α-FLAG) values are used to quantify the relative cell-surface expression of each construct.

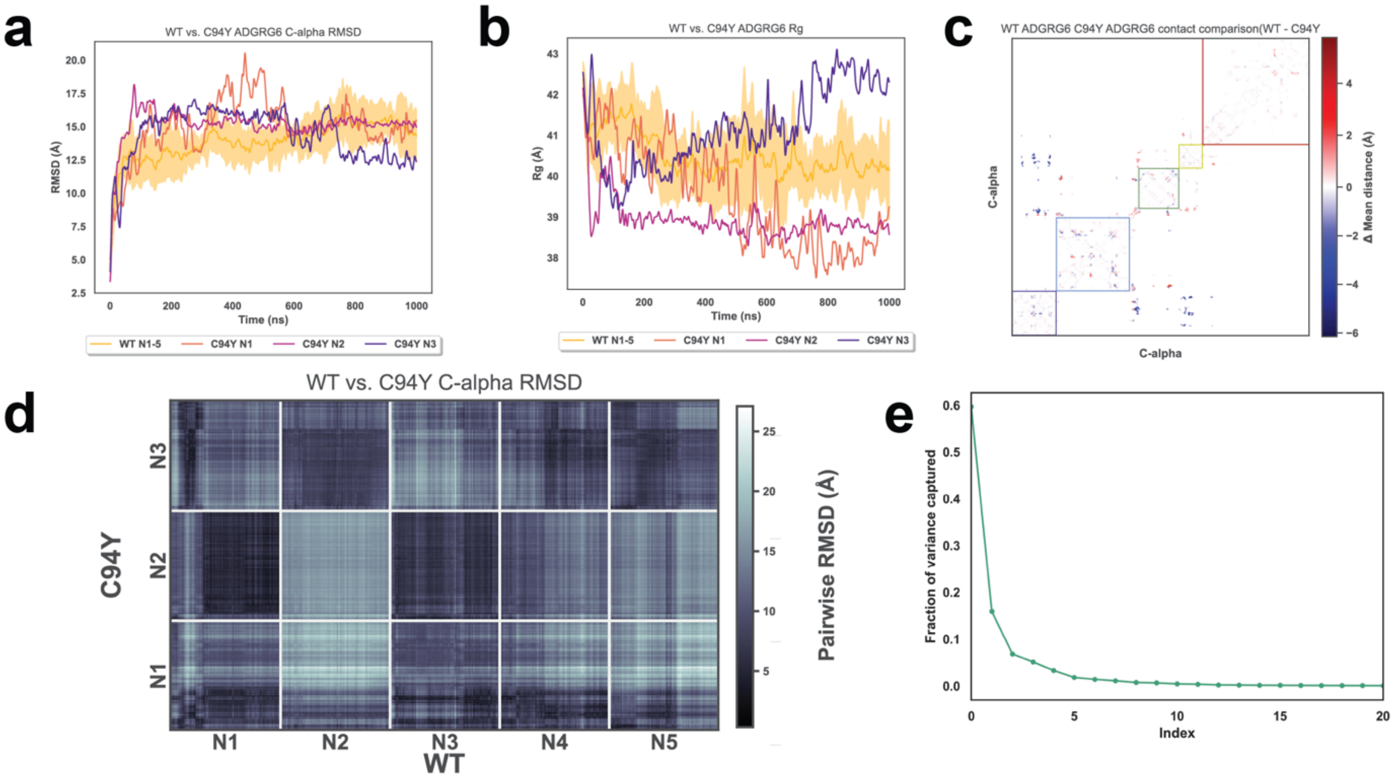


**Fig. S5. Dynamics of the ADGRG6 –ss ECR C94Y simulations. (a):** Cα RMSD of three C94Ysimulations (solid lines) compared to the WT simulations (yellow line and shaded region = mean ± standard deviation). **(b):** $R_g$ comparison between the C94Y and WT simulations. **(c):** Cα–Cα contact comparison between the WT and C94Y simulations. Negative values (blue) represent a greater distance in C94Y than WT, and positive values (red) represent a closer distance in C94Y than WT. **(d):** Cα RMSDs between every structure in the C94Y and WT simulations. **(e):** Scree plot showing the variance of the C94Y simulations captured by each PC. The first 20 PCs are shown for clarity.

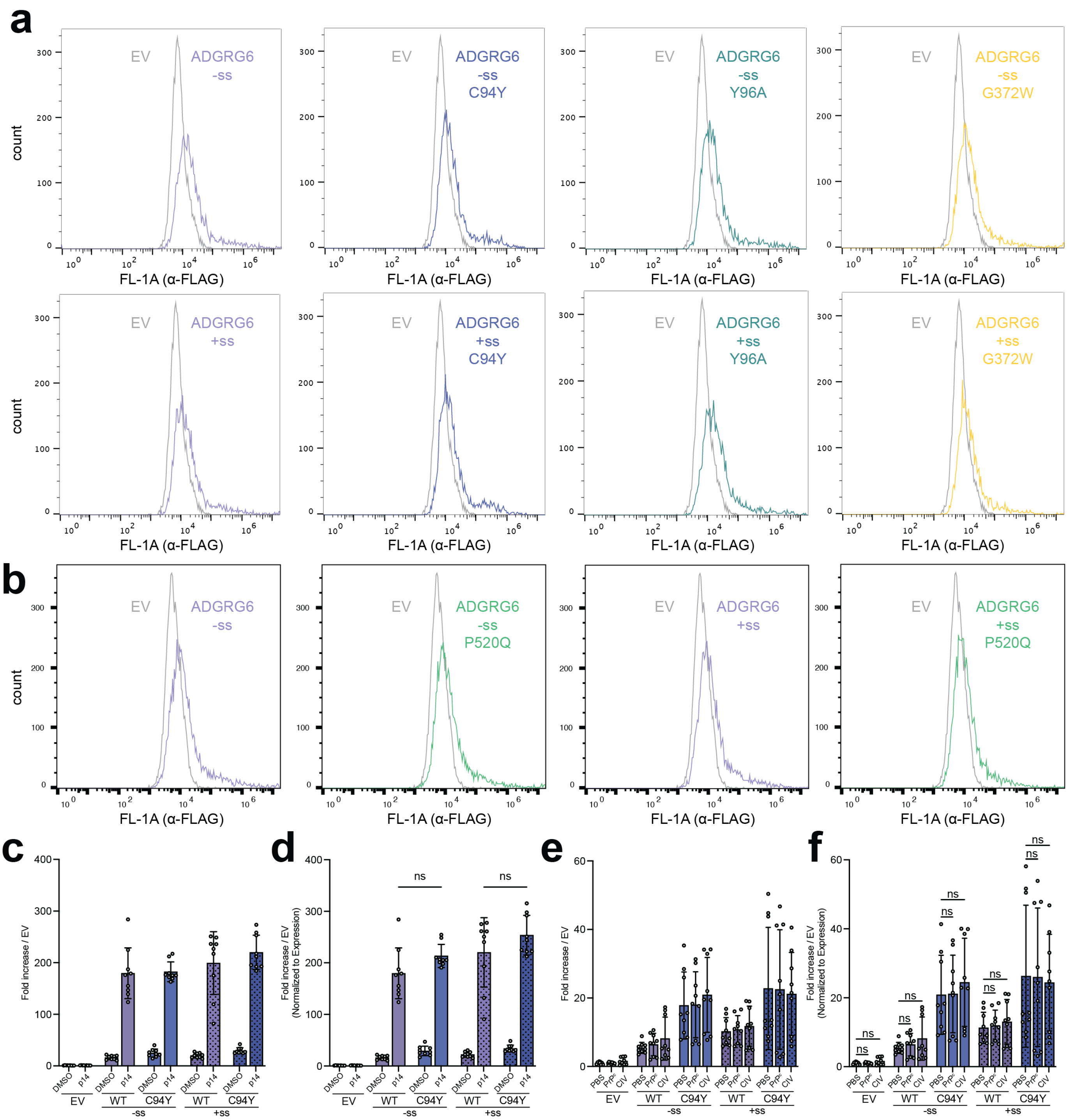


**Fig. S6. Expression levels for ADGRG6 and cancer mutants. (a-b):** Flow cytometry data showing expression levels of the N–terminally FLAG–tagged constructs used for cAMP signaling assays in Figure 6. These are measured using AlexaFluor–488 labeled α–FLAG antibodies. Each plot is overlaid with EV from the same experiment to show background labeling of cells. Median FL-1A (α-FLAG) values are used to quantify the cell-surface expression of each construct. **(c):** Basal cAMP signaling of ADGRG6 constructs shown as fold change over empty vector (EV) in response to addition of Dimethyl Sulfoxide (DMSO) as a loading control or the 14-residue tethered agonist peptide of ADRGR6 (p14). Constructs are grouped by absence (–ss) or presence (+ss) of the ECR splice insert. N=3 independent experiments were performed with three technical replicates each. **(d):** Basal cAMP signaling from (i) represented as fold change over EV normalized to cell-surface expression. ns – not significant; one–way ANOVA test with Tukey's correction for multiple comparisons. Each condition was compared to every other condition, but not all comparisons are shown for clarity. N=3 independent experiments were performed with three replicates each. **(e):** Basal cAMP signaling of ADGRG6 constructs shown as fold change over empty vector (EV) in response toaddition of Phosphate Buffered Saline (PBS) as a loading control, the flexible tail of the cellular prion protein ($PrP^c$), or Collagen IV (CIV). Constructs are grouped by absence (–ss) or presence (+ss) of the ECR splice insert. N=3 independent experiments were performed with three technical replicates each. **(f):** Basal cAMP signaling from (i) represented as fold change over EV normalized to cell-surface expression. ns – not significant; one–way ANOVA test with Tukey's correction for multiple comparisons. Each condition was compared to every other condition, but not all comparisons are shown for clarity. N=3 independent experiments were performed with three replicates each. Bar graphs represent mean values with the standard deviation being shown for error.
